# Multi-Omics Data of Perturbation Studies are Determined by Memory Effects from Subculture

**DOI:** 10.1101/2023.02.13.528316

**Authors:** Patricia Bortel, Gerhard Hagn, Lukas Skos, Andrea Bileck, Verena Paulitschke, Philipp Paulitschke, Lion Gleiter, Thomas Mohr, Christopher Gerner, Samuel M. Meier-Menches

## Abstract

Mass spectrometry-based omics technologies are increasingly used to map drug effects to biological pathways by identifying significant molecular events. Significance is influenced by the effect size and the variation of each molecular parameter. While the former is largely determined by the biological system, the latter can be tuned by the experimental workflow. Here, we unequivocally show that memory effects originating from subculture of colon carcinoma cells before treating with arsenic trioxide exacerbate the variation of multiple omics levels, including eicosadomics, proteomics and phosphoproteomics, without necessarily impacting on effect size. Real-time monitoring of individual samples enables control over subculture homogeneity and improves the median variation >2-fold across omics levels. This considerably facilitated mode of action deconvolution and resulted in a bilevel perturbation network of 321 causal conjectures. Controlling memory effects from subculture revealed key signaling cascades and transcriptional regulatory events that extend the molecular understanding of arsenic trioxide in solid tumors.

## Introduction

Mass spectrometry (MS)-based omics techniques are increasingly employed to characterize drug-induced perturbations because of their hypothesis-generating power.^1^ Perturbation studies are enabled by the identification of molecular events that are mapped to biological processes and the resulting perturbation networks aid in deconvoluting drug modes of action (MoA).^1-3^ Additionally, integrating multi-omics data allows to infer interaction networks that are not accessible by single omics levels^4-6^ and is a powerful tool for causal reasoning.^7,8^

Importantly, the information content of perturbation networks, *i*.*e*. the number of significant and comprehensible molecular alterations, is determined by experimental and statistical workflows. Omics experiments produce large numbers of variables that are accounted for by stringent significance testing.^9,10^ Significance calculations based on p-values consider the effect size and variation of each molecule,^11^ which are put into context of all detected molecules during an experiment by multiple testing corrections, *e*.*g*. false discovery rates (FDRs).^9^ While the effect size reports the mean value difference of a molecule that is characteristic for the perturbation of a system,^11^ the variation is largely determined by the experimental procedure.^12^ Variation is quantified as the relative standard deviation or coefficient of variation (CV) and serves as a key measure for the precision and thus, reproducibility of an experiment or method.^13^ Indeed, the reproducibility crisis is a central challenge in omics sciences^14-16^ and life sciences in general.^17-21^

Early drug discovery is routinely performed using *in vitro* cell cultures. Perturbation studies are evaluated by the independent analysis of a number of biological replicates of control and drug-treated cells to allow for significance testing.^22^ In this setup, instrumental MS-based variation is typically low,^14,22,23^ while sample preparation methods may have considerable, but predictable effects on experimental variation across omics levels.^24,25^ However, the variation induced by biological replicates in one experiment or by the temporal evolution of cell lines during culture may be large and highlights cell culture conditions as a limiting factor for experimental omics data.^26,27^ For example, we evidenced previously that the batch-dependent differences in eicosanoid content of fetal calf serum in the cell culture medium modulates cellular effector functions.^28^ Others described that the pH of cell culture medium differentially regulates intracellular pathways across omics layers, including transcriptomics, proteomics and metabolomics.^29^ Genetic heterogeneity and instability affect cell phenotypes and drug responses.^27,30^ Moreover, cell lines may respond differently to the same perturbation depending on basal expression,^31^ cellular plasticity^32^ or resistance phenomena, *e*.*g*. the epithelial-mesenchymal transition.^26,33^

Here, we demonstrate that so far elusive memory effects from subculture exacerbate the variation, but not necessarily the effect size, nor the number of identified molecules in multi-omics data, including eicosadomics, proteomics and phosphoproteomics by studying the effects of the anticancer agent arsenic trioxide (ATO)^34,35^ on SW480 colon carcinoma cells. Subculture describes the process of transferring a number of cells from one culture flask to another and represents a necessary routine to expand cell lines.^36^ The memory effects result from the passaging of cells at different phases of growth and impact on the homogeneity of cellular responses to perturbations. This subculture phenomenon is distinct from confluence-dependent treatment effects^37,38^ or phenotypic switches caused by genetic instability after long-term culture.^27,30^ Crucially, control over these memory effects improves variation and enables a detailed drug MoA deconvolution. We show that this control can be achieved by real-time monitoring^39,40^ of each individual sample, which may be implemented as a check-point preceding multi-omics workflows.

## Results

### Experimental Design to Assess Memory Effects from Subculture

Perturbation experiments *in vitro* are preceded by a subculture routine, *i*.*e*. by the subdivision of a proliferative cell population.^36^ Clearly, the adhesion and treatment phases after subculture are influenced by the proliferative state of cells before the subculture. This dependency may manifest in memory effects that impact on multi-omics data of perturbation experiments, but their extent was never quantified. The impact of cellular memory effects after subculture was assessed by performing the same perturbation using either homogeneous or heterogeneous subcultures (Figure 1). First, the homogeneous subculture was generated from SW480 colon carcinoma cells in the Log phase of the growth curve^36^ corresponding to approximately 75% growth area occupied by cells. Two 6-well plates were prepared exclusively from this Log phase. Second, a heterogeneous subculture was generated from SW480 cells in the Lag phase, Log phase and plateau phase in separate cell culture flasks. Cells from the Lag phase were subcultured at 20% growth area occupied, while cells from the plateau phase were subcultured at >95% growth area occupied, the latter reflecting density limitation of cell proliferation (Supplementary Information, Figure 1).^36^

**Figure 01.**
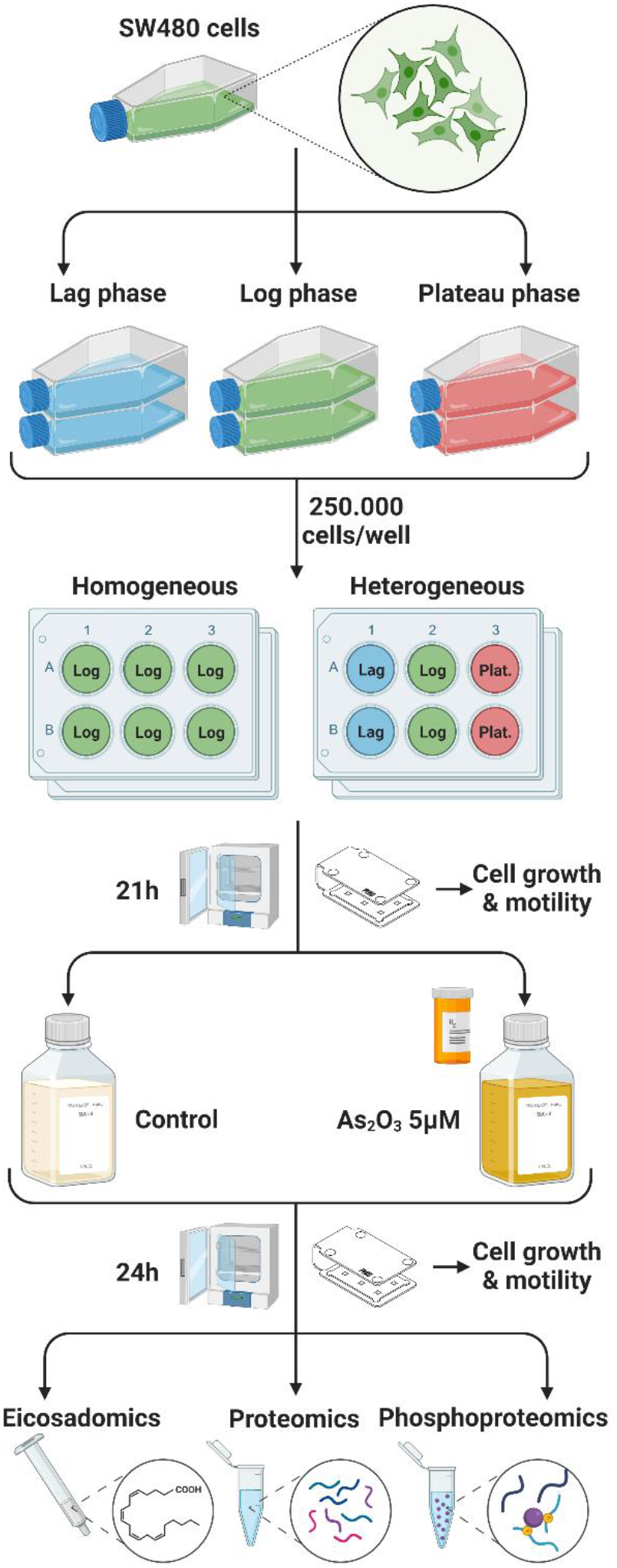
Experimental design to assess the impact of memory effects after subculture on multi-omics data. SW480 colon carcinoma cells were cultured in complete medium and divided into three subcultures, each maintained at different growth phases. The first subculture was maintained in the Lag phase, the second in the Log phase and the third in the plateau phase. Arsenic trioxide treatments (5 μM, 24 h) were performed after homogeneous subculture (cells only from Log phase) or heterogeneous subculture (cells from all three growth phases in duplicates) using always equal cell numbers per well. Cell growth and motility was continuously monitored using a PHIO Cellwatcher M. All samples were processed according to a multi-omics workflow, including oxylipins and fatty acids from supernatants, as well as proteins and phosphoproteins from whole cell lysates.

In this case, two 6-well plates were again prepared. Duplicates from the three different growth phases were seeded in both 6-well plates. In all cases 250’000 cells were seeded per well (Figure 1). The drug treatment was carried out identically after both homogeneous and heterogeneous subcultures. After an adhesion phase, the medium was exchanged and cells were either treated with vehicle control or drug (5 μM) for 24 h. The perturbation was induced with arsenic trioxide (ATO), a clinically approved anticancer agent for the treatment of acute promyelocytic leukemia,^34,35^ which shows a multimodal MoA in cancer cells.^32,41^ Additionally, cell growth and motility was continuously assessed throughout the experiment.

The samples were then processed to enable a multi-omics analysis of each *in vitro* sample adapting a blood plasma analysis procedure established by us.^4^ Here, oxylipins and fatty acids were isolated from supernatants. The protein fraction was obtained from whole cell lysates (WCLs) and was used for global proteomics and phosphoproteomics. WCLs were homogenized for down-stream protein quantification. Oxylipins and fatty acids, as well as global proteomes and phosphoproteomes were analyzed by dedicated liquid-chromatography tandem mass spectrometry (LCMS/MS) methods on different platforms in a data-dependent analysis mode. All reported p-values were multiple testing corrected using FDR calculations and the significantly regulated molecules are listed in the Supplementary Tables 1 and 2. Directed bilevel perturbation networks of the ATO-perturbations after homogeneous and heterogeneous subcultures were calculated from the proteomic and phosphoproteomic data.^7^

### Impact of Subculture Memory Effect on Cell Growth and Motility

Cellular growth and motility was continuously monitored throughout the adhesion and treatment phases using the Cellwatcher M, which combines six microscope units in one device and is compatible with 6-well plates. The memory effects from subculture are clearly illustrated by the growth area occupied by cells. Initially, the SW480 cells were seeded at around 10–15% growth area occupied. The increase in growth area occupied over the first day relates to cell spreading and proliferation. Crucially, although seeded at equal cell numbers, cell growth after homogeneous subculture is much more uniform among the six individual replicates (Figure 2A) compared to cell growth after heterogenous subculture (Figure 2B). ATO-treatment led to an immediate growth inhibition irrespective of subculture condition. At the last analysis before harvesting, the control cells showed 89±5% (CV = 6%) and 84±11% (CV = 13%) growth area occupied after homogeneous and heterogeneous subculture, respectively. The treatment clearly reduced cell growth to 57±4% (CV = 7%) after homogeneous subculture, while this effect was less pronounced after heterogeneous subculture showing 77±18% (CV = 23%) growth area occupied.

**Figure 02.**
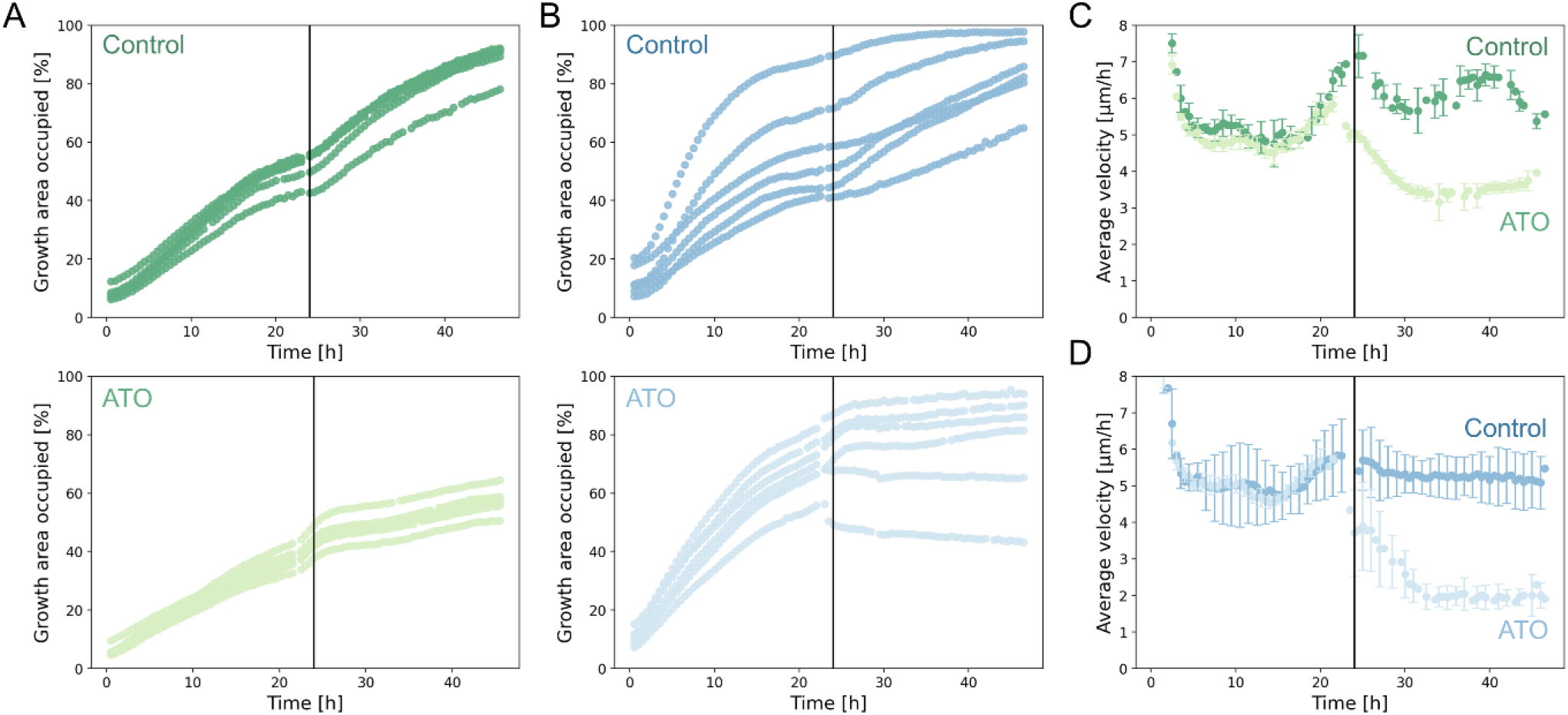
Homogeneous subculture improves variation of cell growth and motility compared to heterogeneous subculture. Cell growth and motility was continuously monitored over the entire adhesion and treatment phases after subculture. Medium was replaced after 24 h either containing vehicle control or arsenic trioxide (ATO, 5 μM) and incubated for further 24 h. The six individual wells were monitored after homogeneous subculture (**A**, green) and after heterogeneous subculture (**B**, blue). Treatment with ATO reduced cell growth independently of subculture conditions (**A**, light green; **B**, light blue). Similarly, cell motility was reduced upon treatment with ATO independently of homogeneous (**C**, green) and heterogeneous (**D**, blue) subcultures, but variation increased in the latter.

Transformed cells typically display random migration patterns, which can be determined by means of time-lapse microscopy. Cell motility depends on the cell density in the plate, but also on drug treatments. At the last analysis before harvesting, the control cells displayed a mean cell motility 5.6±0.2 μm·h^−1^and 5.5±0.7 μm·h^−1^after homogeneous and heterogenous subculture, respectively. ATO-treated cells showed a reduced motility of 4.0±0.2 μm·h^−1^ and 1.9±0.2 μm·h^−1^ after homogeneous and heterogenous subcultures, respectively (Figure 2C and 2D; Supplementary Information, Figure 2). The drug treatment led to a substantial decrease in cell motility that complements growth inhibition irrespective of the subculture condition. Importantly, the homogeneous subculture delivered an improved variation of the biological replicates compared to the heterogeneous subculture. The controls from homogeneous subculture displayed a CV of 4% of cell motility at the last time point before harvesting, while the controls from heterogeneous subculture showed a CV of 12%. The impact of subculture was similar for the treated cells, where the homogeneous and heterogeneous subcultures yielded CVs of 5% and 10%, respectively.

Overall, the phenotypic parameters of cell growth and motility evidenced at least two-fold improved variation after homogeneous compared to heterogeneous subculture.

### Impact of Subculture Memory Effect on Oxylipins and Fatty Acids

Oxylipins and fatty acids were isolated from the supernatants of control and ATO-treated SW480 cancer cells. A total of 87 oxylipins and fatty acids were identified after both homogeneous and heterogeneous subculture, including 27 eicosanoids. Of the 87 identified oxylipins and fatty acids, 31 were significantly upregulated upon treatment with ATO after homogeneous subculture, while only 10 were upregulated after heterogeneous subculture. None were significantly downregulated. The latter 10 lipid species were all found in the data after homogeneous subculture as well (Figure 3A).

**Figure 03.**
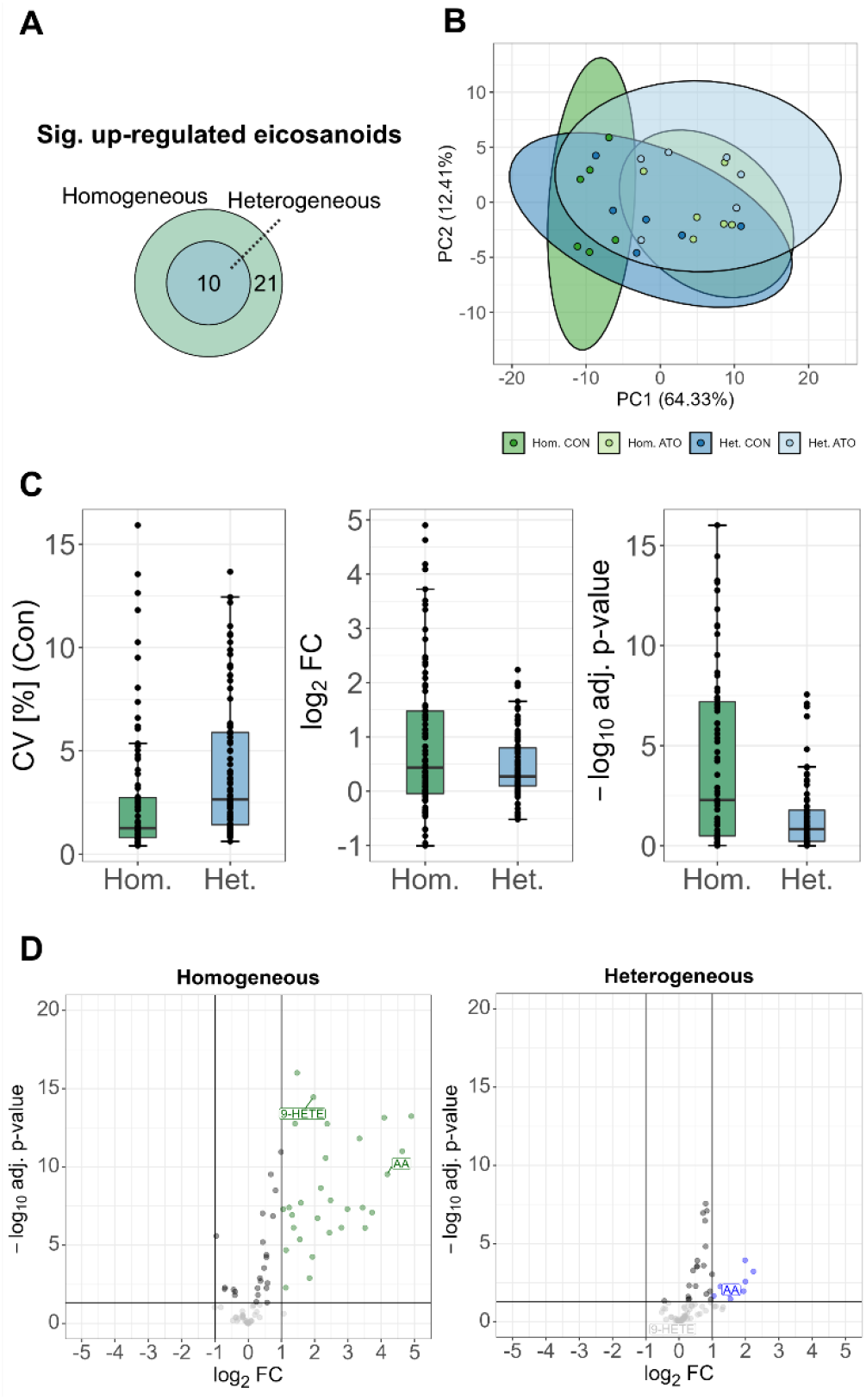
Oxylipin and fatty acid data after homogeneous subculture shows an improved variation compared to data obtained after heterogeneous subculture. (**A**) The Venn diagram illustrates the shared and specific upregulated eicosanoids upon treatment of SW480 cancer cells with arsenic trioxide (ATO). (**B**) Principal component analysis (PCA) of normalized area under the curve (nAUC)-intensity values of identified oxylipins and fatty acids in control and ATOtreated SW480 cells after homogeneous (green) and heterogeneous (blue) subcultures. The first two principal components separate controls and ATO-treatments after homogeneous, but not after heterogeneous subculture. Ellipses represent 95% confidence intervals. (**C**) Boxplots of the control samples displaying coefficients of variance (CV) of nAUC-intensities of oxylipins and fatty acids among six biological replicates after homogeneous (green) and heterogeneous (blue) subcultures. Also, the log_2_-fold-changes (FC) of the perturbation of nAUC-intensities was plotted as boxplots after homogeneous and heterogeneous subcultures, as well as the associated multiple testing corrected p-values. (**D**) Volcano plots showing the ATO-induced perturbations after homogeneous (green) and heterogeneous (blue) subcultures. Significant regulations after multiple testing correction are colored.

Principal component analysis (PCA) showed an improved variation of eicosadomic profiles among replicates after homogeneous subculture compared to heterogeneous subculture (Figure 3B). Accordingly, control and ATO-treated samples were clearly separated after the homogeneous subculture, while samples corresponding to the respective conditions were found more dispersed after the heterogeneous subculture and overlapped considerably.

The median CV of normalized intensity values among replicates of the control group was 1% after homogeneous subculture and 3% after heterogeneous subculture (Figure 3C). For this compound class, ATO treatment affected the amplitude of the fold-change distributions. The data after homogeneous subculture was more dynamic compared to the treatment after heterogeneous subculture. In contrast, the median log_2_ fold-changes were similar after the homogeneous (0.43) and heterogeneous subcultures (0.27). As a consequence, the data set after homogeneous subculture displayed improved adjusted (adj.) p-values compared to the heterogeneous subculture, which is highlighted in median –log_10_ adj. p-values of 2.3 and 0.8, respectively.

Upon ATO treatment, arachidonic acid (AA) was among the shared upregulated lipid species after both subcultures (Figure 3D). AA is the precursor of most eicosanoids^42^ and it was found significantly upregulated 18– (CV = 3%, –log_10_ -adj. p-value = 9.5) and 4fold (CV = 8%, –log_10_ -adj. p-value = 2.6) after homogeneous and heterogeneous subcultures, respectively. In contrast, important signaling mediators of AA were only significantly upregulated after homogeneous subculture. For example, the non-enzymatically formed 9-hydroxyeicosatetraenoic acid (9-HETE), which is a marker for oxidative stress,^43^ was significantly upregulated only after homogeneous subculture (FC = 3.9, – log_10_ -adj. p-value = 14.5), but not after heterogeneous subculture (FC = 1.2, −log_10_ -adj. p-value = 0.8). In addition, the epoxy-eicosatrienoic acids (EETs) are cytochrome P450 oxidation products of AA with anti-apoptotic and anti-oxidative properties.^44^ The identified 5(6)EET and 14(15)-EET were significantly upregulated 10.8– (−log_10_ -adj. p-value = 7.4) and 4.5-fold (−log_10_ adj. p-value = 8.7) after homogeneous subculture. After heterogeneous subculture, 5(6)-EET showed a fold-change of 2.1 (–log_10_ -adj. p-value = 1) and 14(15)-EET a fold-change of 1.1 (–log_10_ -adj. p-value = 0.1).

Thus, memory effects from subculture impact on global oxylipin and fatty acid profiles. Homogeneous subculture led to >2-fold improved CV and improved p-values compared to heterogeneous subculture and resulted in a >3-fold increase in the number of significant regulations.

### Impact of Subculture Memory Effect on Proteomics

The proteomic analysis of whole cell lysates resulted in the identification of a total of 4554 and 4445 proteins after the homogeneous and the heterogeneous subculture, respectively. A total of 4291 proteins were identified after both the homogeneous and heterogeneous subcultures, while 263 and 154 proteins were exclusively identified after the respective subcultures (Figure 4A). Of the 343 significantly upregulated proteins upon treatment with ATO, 106 (31%) were upregulated after both subculture conditions, while approximately the same numbers were upregulated after one specific subculture condition. A more pronounced contrast was evidenced for the down-regulated proteins, where 83 (12%) proteins of a total of 681 downregulated proteins were shared between the two subculture conditions. The homogeneous subculture resulted in 458 uniquely down-regulated proteins compared to 140 after heterogeneous subculture.

**Figure 04.**
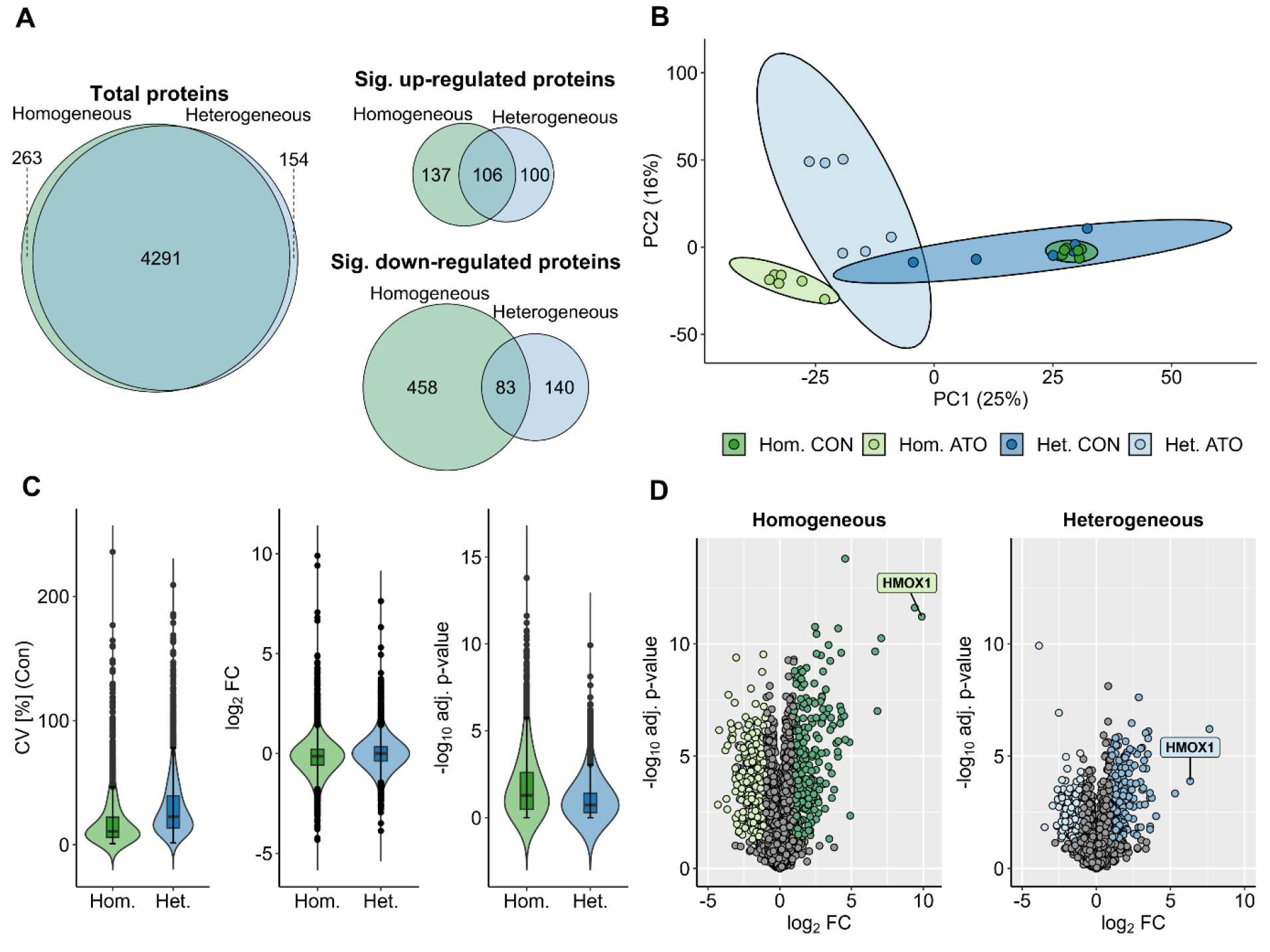
Homogeneous subculture improves variation of proteomics data compared to heterogeneous subculture. (**A**) The Venn diagrams illustrate shared protein identifications upon treatment with arsenic trioxide (ATO) after homogeneous and heterogeneous subcultures, as well as shared significantly upand down-regulated proteins (adj. p-value ≤0.05, fold change ≥1 or ≤1). (**B**) Principal component analysis (PCA) of proteome profiles in control and ATO-treated SW480 cells after homogeneous (green) and heterogeneous (blue) subculture. The first two principal components separate homogeneous controls from ATO-treatments, while this was not observed after heterogeneous subculture. Ellipses represent 95% confidence intervals. (**C**) Violin plots of the control samples displaying the coefficients of variance (CV) of LFQ-intensities among six replicates after homogeneous (green) and heterogeneous (blue) subcultures. The log_2_ -fold-changes (FC) after homogeneous and heterogeneous subcultures are displayed as violin plots, as well as the associated multiple testing corrected p-values. (**D**) Volcano plots showing the global ATO-induced perturbations after homogeneous (green) and heterogeneous (blue) subcultures on the protein level. Multiple testing corrected significant protein regulations are colored.

The PCA showed an improved variation of replicates after the homogeneous compared to the heterogeneous subculture (Figure 4B). Accordingly, control and ATO-treated samples were clearly separated after the homogeneous subculture, while samples after the heterogeneous subculture exhibited a partial overlap.

The median CV of protein LFQ intensity values among replicates of the control group was 11% after the homogeneous subculture and 23% after the heterogeneous subculture (Figure 4C). Regarding protein regulations upon ATO treatment, the median log_2_ foldchanges were similar, corresponding to −0.13 and 0.01 after homogeneous and heterogeneous subculture, respectively. The median −log_10_ -adj. p-value accounted for 1.3 and 0.7 after the homogeneous and heterogeneous subcultures, respectively.

Upon ATO treatment, 243 proteins were significantly upregulated and 541 proteins significantly downregulated after the homogeneous subculture. After heterogeneous subculture, 206 and 223 proteins were significantly up- and down-regulated upon ATO treatment, respectively. The impact of memory effects from subculture was also evidenced at the single protein level. For example, heme oxygenase 1 (HMOX1), which is involved in protection of cells against oxidative stress and apoptosis,^45^ was among the most strongly upregulated proteins after both the homogeneous and the heterogeneous subcultures, exhibiting log_2_ foldchanges of 9.9 and 6.33, respectively (Figure 4D). Yet, the associated –log_10_ -adj. p-value was considerably reduced from 11.2 after homogeneous to 3.9 after heterogeneous subculture.

For proteomics, a homogeneous subculture improved the experimental variation 2-fold and also improves the overall p-values. However, the two subculture conditions led to sparse overlap in the perturbation experiment.

### Impact of Subculture Memory Effect on Phosphoproteomics

The phosphoproteomic analysis resulted in the identification of 2278 and 2082 class I phosphosites with localization probability ≥75% after the homogeneous and the heterogeneous subculture, respectively. The number of phosphosites identified after both subcultures was 1812, corresponding to 71% of the total phosphosites (Figure 5A). A total of 466 and 270 phosphosites were uniquely identified after the homogeneous and heterogeneous subcultures, respectively. Of the 618 significantly upregulated phosphosites only 32 (5%) were shared between the two subculture conditions, while 538 phosphosites were uniquely upregulated after the homogeneous subculture and 48 were uniquely upregulated after the heterogeneous subculture. A similar amount corresponding to 29 (9%) of a total of 312 phosphosites were significantly down-regulated after both subculture conditions, while 134 phosphosites were uniquely down-regulated after the homogeneous subculture and 149 were uniquely downregulated after the heterogeneous subculture.

**Figure 05.**
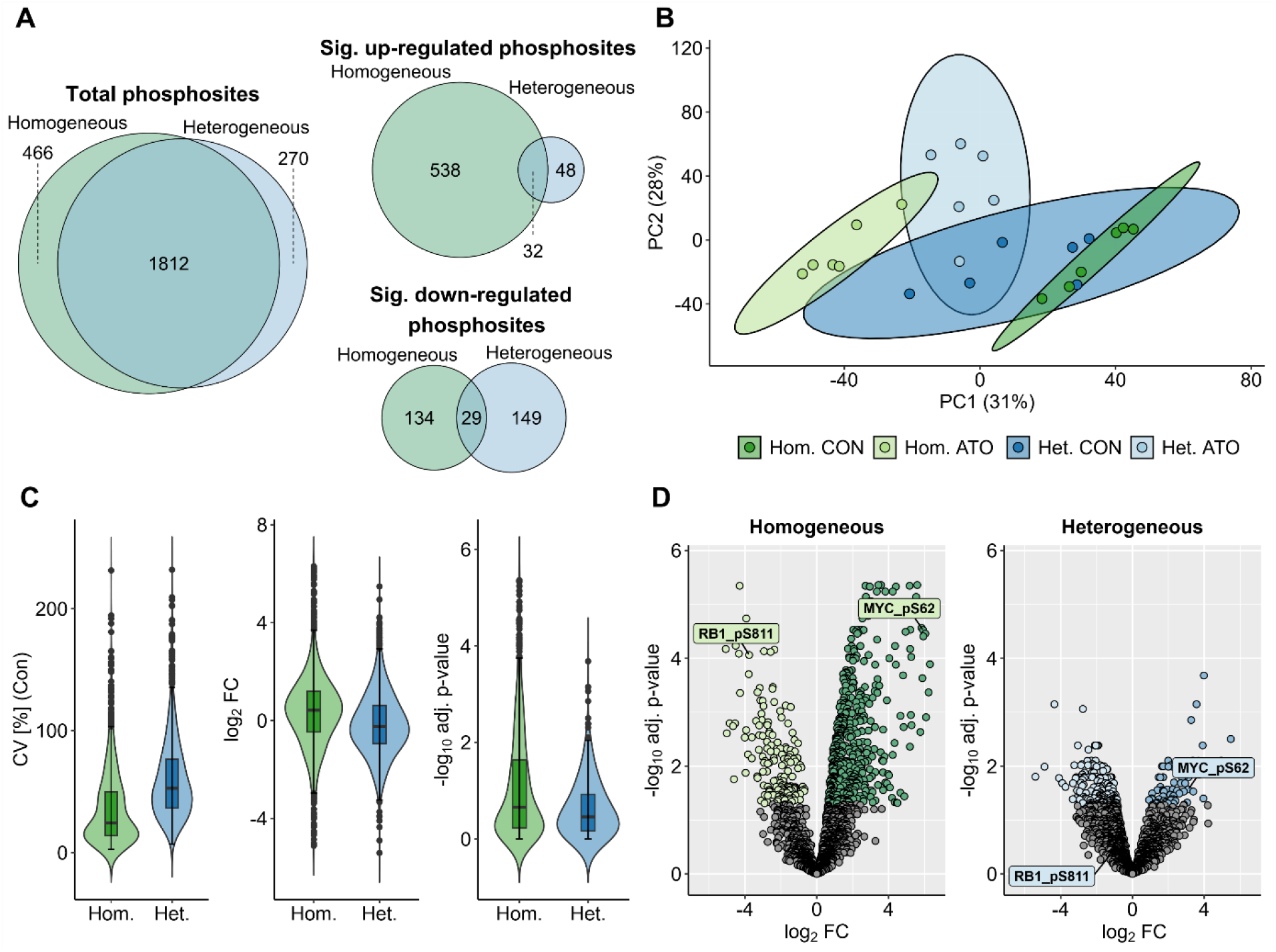
Homogeneous subculture improves variation of phosphoproteomic data compared to heterogeneous subculture. (**A**) The Venn diagrams illustrate shared phosphosite identifications upon treatment with arsenic trioxide (ATO) after homogeneous and heterogeneous subcultures, as well as shared significantly upand downregulated phosphosites (adj. pvalue ≤0.05, fold change ≥1 or ≤1). (**B**) Principal component analysis (PCA) of phosphosite profiles of control and ATO-treated SW480 cells after homogeneous (green) and heterogeneous (blue) subcultures. The first two principal components separate homogeneous controls from ATO-treatments, while this was not the observed after heterogeneous subculture. Ellipses represent 95% confidence intervals. (**C**) Violin plots of the control samples displaying the coefficients of variance (CV) of LFQ-intensities of phosphosites among six replicates after homogeneous (green) and heterogeneous (blue) subculture. Also, the log_2_ -foldchanges (FC) of phosphosite LFQ-intensities were plotted as violin plots, as well as the associated multiple testing corrected pvalues. (**D**) Volcano plots showing the ATO-induced perturbation after homogeneous (green) and heterogeneous (blue) subcultures on the phosphosite level. Multiple testing corrected significant protein regulations are colored.

The PCA of the phosphoproteomics data showed that replicates after heterogeneous subculture were more dispersed compared to the replicates after homogeneous subculture (Figure 5B). Similarly, ATO-treated samples were clearly separated from the control samples after homogeneous subculture, whereas samples after heterogeneous subculture overlapped considerably.

The median CV of phosphosite LFQ intensity values among the replicates of the control group was 24% after homogeneous subculture and 53% after heterogeneous subculture (Figure 5C). While the dynamics of the fold-change distributions remained similar, the median log_2_ fold-change of phosphosites upon ATO treatment was 0.42 and –0.24 after homogeneous and heterogeneous subculture, respectively. The median – log_10_ -adj. p-value accounted for 0.7 and 0.5 after the homogeneous and heterogeneous subculture, respectively.

Memory effects from subculture were also evidenced at the single phosphopeptide level (Figure 5D). For example, phosphorylation of the transcription factor MYC at S62, a central event of the cellular response to ATO,^46^ was strongly induced after both subculture conditions. However, this phosphorylation event was much more pronounced after homogeneous (log_2_ foldchange 5.92, –log_10_ -adj. p-value 4.5) compared to heterogeneous subculture (log_2_ fold-change 3.12, –log_10_ adj. p-value 1.6). Moreover, the transcription factor RB1 was found dephosphorylated at S811 upon treatment with ATO. While this dephosphorylation event was significant after homogeneous subculture, it was not found significant after heterogeneous subculture.

The significantly upregulated phosphosites were mapped to potential kinases *via* the kinase substrate scoring function of PhosR^47^ based on their dynamic phosphorylation profiles and kinase recognition motifs. This combined score includes a minimum of 5 sequences used for compiling the motif for each kinase and minimum of one phosphosite for compiling phosphorylation profiles for each kinase. A heatmap of the combined scores retrieved for the top three phosphosites with the highest combined scores of all evaluated kinases is shown in Supplementary Information, Figure 3. The mapping of significantly upregulated phosphosites after the homogeneous subculture resulted in the annotation of 40 kinases to their corresponding top 3 scoring phosphosites (Supplementary Information, Figure 3A), whereas 19 kinase-substrate pairs were determined after the heterogeneous subculture (Supplementary Information, Figure 3B).

A strong impact of memory effects from subculture was observed at the level of the phosphoproteome. Again, the variation improved 2-fold after homogeneous compared to heterogeneous subculture, while the p-value distribution also improved for the former. Although a similar number of phosphosites were identified there was scarce overlap between significant phosphosite regulations, which was found <10%.

### Impact of Subculture Memory Effect on the Perturbation Network

Proteomic and phosphoproteomic analyses were integrated into a bilevel causal perturbation network using CausalPath.^7^ After homogeneous subculture, a total of 321 causal conjectures were identified (Figure 6A), while 58 causal conjectures were observed after heterogeneous subculture (Figure 6B). The majority of the causal conjectures found in the perturbation network after heterogeneous subculture were also found in the one after homogeneous subculture. Of these shared conjectures, the directions of individual protein regulation and (de-)phosphorylation events were also similar. ATO is known to exert pleiotropic effects by inducing multiple signaling cascades and affecting numerous cellular functions.^32,46^ Notable contributors to the MoA beyond PML-RARα degradation include the induction of reactive oxygen species (ROS), protein kinase signaling, heat shock proteins, apoptosis, as well as the disruption of the cell cycle.^32,46^

**Figure 06.**
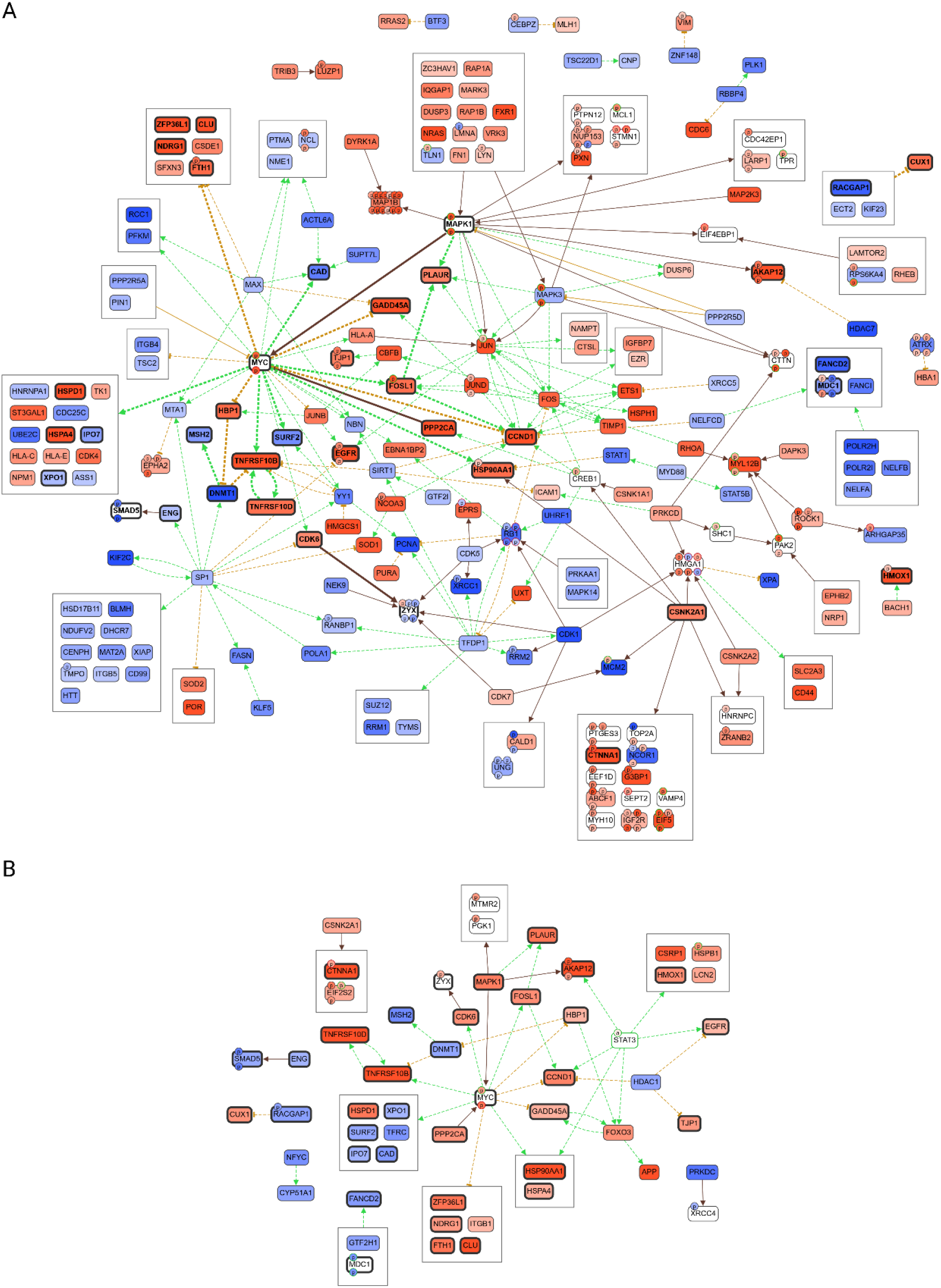
Bilevel CausalPath analysis using global proteomic and phosphoproteomic data of arsenic trioxide (ATO)induced perturbation networks after homogeneous (A) and heterogeneous (B) subcultures. The homogeneous and heterogeneous subcultures yielded 321 and 58 causal conjectures, respectively. The memory effect after homogeneous subculture improves experimental variation and enhances the information content of the resulting perturbation network for the ATO-treatment on SW480 cancer cells. Red represents upregulated gene expression products or phosphorylation events. Blue represents down-regulated gene expression products or dephosphorylation events. Dashed lines represent gene expression changes (green = positive correlation of expression, orange = negative correlation of expression. Continuous-line brown arrows represent phosphorylation events. Phosphosites may be activating (green frame) or inactivating (red frame). STAT3 was annotated (“a”) by the CausalPath algorithm due to the concerted upregulation of gene expression products in the perturbation network after heterogeneous subculture (**B**). Bold lines and boxes in (**A**) reflect those causal conjectures that were also identified in (**B**). *A high-resolution image of the perturbation network is provided as a supplementary figure*.

Both perturbation networks featured robust ATO-induced expression changes and phosphorylation events that are coordinated by the transcription factor MYC (Figure 6).^46^ ATO treatment activated MAPK1, which directly phosphorylated MYC^48^ and this is paralleled by EGFR activation.^49^ Arsenic is known to decrease transcription by direct effects on the transcription factors SP1 and MYC,^50^ and this seems to be further supported by the upregulation of ZFP36L1,^51^ which mediates RNA decay. Moreover, a reduced transcriptional activity of MYC led to the upregulation of the transcriptional repressor HBP1, which plays a role in cell cycle regulation by down-regulating the S-phase associated DNA methyltransferase DNMT1.^52^ Similarly, ATO induced the expression of GADD45A^53^ and cyclin D1 (CCND1),^50^ which are regulators of the G2/M and G1/S transitions, respectively. DNA damage repair signatures were characterized by the down-regulation of MSH2^54^ and FANCD2. Next to initiating MAP signal transduction pathways, ATO induced several heat shock proteins, including HSP90AA1, HSPD1 and HSPA4. HSP90AA1 (HSP90) is the most abundant chaperone and seems to be inhibited by arsenite^55^ and HSPD1 (HSP60) was shown to be a direct target of ATO.^56^ Additionally, as part of the stress response, HMOX1^32^ and NDRG1 were also found upregulated upon ATO treatment in both perturbation networks, the latter being necessary for p53-dependent apoptosis.^57^ ATO also induced the death receptors TNFRSF10B and TNFRSF10D.^58^ The upregulation of the casein kinase II subunit alpha (CSNK2A1)^59^ was observed and led to the phosphorylation of a number of substrates in both perturbation networks, *e*.*g*. a deactivating phosphorylation on catenin alpha-1 (CTNNA1), which affects the actin network and cadherin-cell adhesion properties. Accordingly, the cytoskeleton was suggested as a mediator of the MoA of ATO.^46^ These effects illustrate the known intracellular effects of ATO.^32,46^

In contrast to these similarities, the perturbation networks also showed specific responses to the ATO treatment depending on the subculture condition. The perturbation network after homogeneous subculture yielded numerous additional causal conjectures compared to the one after heterogeneous subculture. This enhanced the information density in the network and highlighted additional transcription factor hubs and signaling events (Figure 6A). For example, the transcription factor SP1^60^ was down-regulated in addition to MYC. SP1 coordinated the down-regulation of 17 gene-expression products, including DNMT1, and the up-regulation of SOD1, SOD2,^61^ and EGFR. This perturbation network also displayed activating phosphorylations on EGFR next to its upregulation.^49^ Additional down-regulated transcription factors not yet implicated in the MoA of ATO included MAX, TFDP1 and RB1, the latter featured decreased phosphosite occupancy.^62^ RB1 is a tumor suppressor protein that is tightly linked to cell cycle regulation^63^ and connected to CDK1 and CDK5 *via* phosphorylation events. Moreover, arsenite is known to activate the transcription factor AP-1 by increasing the expression of the mitogenic components JUN and FOS.^64^ Those were exclusively observed in the perturbation network after homogeneous subculture and were upregulated in contrast to the other transcription factors. Phosphorylation of JUN was mediated by MAPK1 and MAPK3, which themselves were detected containing activating phosphorylations. Activated MAPK1 also led to the hyperphosphorylation of the microtubulin-binding protein MAP1B that may reflect cytoskeletal adaptions.^46^ Upregulated AP-1 also induced RHOA signaling,^32^ which was exclusively observed in this perturbation network.

The perturbation network after heterogeneous subculture featured only limited specific effects, for example STAT3 as a second hub of controlling gene expression. The effect of arsenic on STAT3 seems to be highly context-dependent.^65,66^ However, this transcription factor was not detected in the data set and was annotated by the CausalPath algorithm based on the upregulation of a number of down-stream gene expression products (Figure 6B). A further down-regulated DNA damage signature including PRKDC and the concomitant dephosphorylation of XRCC4 was observed, which are involved in non-homologous end joining.

Consequently, homogeneous subculture facilitates MoA deconvolution in perturbation studies and yields a higher information density for pathway discovery compared to perturbations after heterogeneous subculture.

## Discussion

MS-based omics techniques are valuable tools for drug discovery.^1,3^ They allow to construct perturbation networks to aid in MoA deconvolution, which may be additionally facilitated by multi-omics combinations,^6,67^ increasing the number of perturbations^2^ or by attempts of causal reasoning.^7^ The quality of the perturbation network is strongly determined by experimental and sample processing workflows that are governed by an overall variation, expressed as CV, which defines the experimental reproducibility. The impact of cell culture conditions on cellular functions and the reproducibility of omics data is a current topic of investigation.^15,27,28^Approaches to account for the genetic evolution of cells in culture *via* cellular force tracking exist and provide quantitative data.^68^ Here, we demonstrate that memory effects of cancer cells from subculture routines directly impact on the variation of multi-omics data and the resulting perturbation network. It may be noted that these effects were obtained irrespective of the statistical evaluations, *i*.*e*. linear modelling using BenjaminiHochberg^69^ or Perseus using permutation-based FDR.^70^

In this study, two different subculture conditions were established to assess the memory effect of SW480 colon cancer cells on ATO-induced perturbations on multiple omics levels (Figure 1). The homogeneous subculture consisting of cells exclusively from the Log phase of cell growth while the heterogeneous subculture was generated using cells from different phases of cell growth, including Lag phase, Log phase and plateau phase. ATO is an intriguing anticancer metalloid that cures patients suffering from acute promyelocytic leukaemia in combination with all-*trans* retinoic acid.^71^ Its clinically relevant MoA includes the degradation of the oncogenic PML-RARα fusion protein that relieves the differentiation blockade of the premature cancerous blasts.^34^ In solid tumors, however, ATO was shown to exert pleiotropic effects based on the induction of ROS, heat shock proteins, kinase signaling and apoptosis.^46^

Although the subculture conditions affected the number of identified molecules on the eicosadomic, proteomic and phosphoproteomic level only to a minor degree (Figures 3A–5A), they considerably affected the number of significantly regulated molecules in favor of the treatment after homogeneous subculture on all levels up to 7-fold, as exemplified by the upregulated phosphosites (Figure 5A). The molecular profiles of the samples obtained after homogeneous subculture were also more uniform on all omics levels and irrespective of control or treatment conditions, compared to samples after heterogeneous subculture. This resulted in an improved group separation in the PCA plots (Figures 3B–5B). Therefore, the homogeneous subculture yielded higher significance values compared to the heterogeneous subculture on all omics levels (Figures 3C–5C). Since the overall fold-change distributions were only affected to a minor degree, this effect can be mainly attributed to a >2-fold improved CV after homogeneous subculture. The volcano plots also clearly illustrate this point at the y-axes (Figures 3D– 5D).

The larger number of significantly regulated molecules after homogeneous subculture resulted in an enhanced information content for delineating MoA. For example, ATO treatment after homogeneous subculture yielded important signaling eicosanoids that were not significantly regulated after heterogeneous subculture. Among others, we found 9-HETE, a marker for oxidative stress,^43^ which is formed non-enzymatically from AA or the P450-products 5(6)-EET and 14(15)-EET, which exhibit anti-apoptotic and anti-oxidative effects.^44^ The known induction of ROS by ATO can thus be followed in the supernatant only after homogeneous subculture.

Although the overlap of significantly regulated proteins (<30%) and phosphosites (<10%) upon ATO treatment was low, the obtained bilevel perturbation networks from proteomic and phosphoproteomic data featured clear similarities. Here again, the increase in the number of significant regulations after homogeneous subculture precipitated in a higher information density. ATO-treatment after homogeneous subculture generated 321 causal conjectures, while the treatment after heterogeneous subculture yielded 58 causal conjectures. Both perturbation networks confirmed essentially the multi-modal MoA of ATO in solid tumors (Figure 6).^46^ In short, we observed the induction of HMOX1,^32^ as well as the heat shock proteins HSP90AA1,^55^ HSPD1^56^ and HSPA4, which indicate the generation of ROS by ATO. ATO treatment was found to activate MAPK1.^46^ This in turn led to phosphorylations on the transcription factor MYC,^48^ which is an important regulator of cell fate and directly controls heat shock protein expression. Phosphorylation of MYC at serine-62^72^ was confirmed as an important initial effect of ATO and was observed after both subculture conditions (Figure 6; Supplementary Information, Figure 3). The impact of ATO on the cell cycle was evidenced by the induced expression of GADD45A^53^ and CCND1,^50^ which are regulators of cell cycle transitions. Apoptotic signals were not observed on the protein levels because we used sub-cytotoxic concentrations of ATO.^32^

Despite the possibility to delineate the expected MoA of ATO after heterogeneous subculture, the homogeneous subculture provided clearly more information. This can be exemplified by the observation of the upregulated transcription factors SP1^60^ and AP1^46,64^ as hubs for mediating ATO-induced gene expression changes, and by numerous cell cycle regulators (Figure 6A). The perturbation network also revealed so far unprecedented effects of ATO, including the downregulation of the transcription factors Max, TFDP1 and RB1. Interestingly, RB1 is directly involved in the regulation of the cell cycle^63^ and featured four dephosphorylation events.^62^ Homogeneous subculture improves the hypothesis-generating power of omics technologies for drug discovery.

A number of cellular pathways affected by the ATO-treatment were not identified directly in the perturbation network, but only *via* manual data interpretation. Among these was the induction of the cytoprotective Nrf2-mediated response against oxidative stress, which was previously shown as a relevant resistance pathway against ATO.^73^ Several canonical Nrf2-target genes were induced upon drug treatment after both subculture conditions, including SQSTM1, SRXN1, GCLM, as well as the ferritin components FTL and FTH1, in additions to the already mentioned HMOX1 (Supplementary Information, Figure 4A). Further Nrf2target genes were observed exclusively after homogeneous subculture, including TXNRD1, GSTM4 and the oxidative stress-induced growth inhibitor 1 (OSGIN1), which may be an indicator for the apparent stress-induced cell cycle arrest. Oxidative stress may eventually lead to DNA damage. However, an ATO-induced DNA damage response was not observed. Oxidative stress on DNA may have been limited by the action of abasic site processing protein HMCES, an ancient DNA lesion recognition protein preserving genome integrity.^74^ This suicide enzyme was found stably expressed in most control cells independently of subculture, but disappeared in virtually all ATO-treated cells (Supplementary Information, Figure 4B).

The impact of subculture conditions was also observed at the phenotypic level. Real-time monitoring of cell growth and motility paralleled the findings of the multi-omics data. Although cell growth and motility were similarly inhibited after both subculture conditions upon ATO treatment, the variation of controls and treated samples were consistently >2-fold improved after homogeneous subculture. Since the improvement of variation at the phenotypic level is comparable to that at the molecular level, real-time monitoring may serve as a quality check-point before performing elaborate multi-omics workflows.

In summary, while confluence dependencies and phenotypic switches in cell culture are known to impact on the quality of omics experiments, this study quantifies the influence of memory effects from subculture on multi-omics data and perturbation networks. A clearly improved experimental variation on eicosadomic, proteomic and phosphoproteomic levels was observed after homogeneous compared to heterogeneous subculture. This increased the number of significantly regulated molecules and consequently enhanced the information density of the perturbation network. Real-time monitoring of cell growth and motility allows to monitor subculture homogeneity and may be implemented as a check-point before performing elaborate multi-omics workflows. The control of memory effects from subculture is key to maximize MoA deconvolution for omics-based drug discovery.

## Methods

### Cell Culture

#### Culture conditions

The colon carcinoma cell line SW480 was kindly provided by Michael Jakupec (Department of Inorganic Chemistry, University of Vienna, Austria). SW480 cells were cultured in Eagle’s minimum essential medium (EMEM; Sigma–Aldrich, USA) supplemented with 10% FCS, 1% penicillin/streptomycin, 2 mM l-Glutamine, 0.1 mM non-essential amino acids and 1 mM sodium pyruvate (all Gibco, USA). The cells were grown in a humidified atmosphere with 5% CO_2_ at 37°C.

#### Treatment

SW480 cancer cells were cultured in T75 flasks for adherent cell culture (Sarstedt GmbH, Nümbrecht, Germany). Prior to the drug treatment, three distinct subculture conditions were generated containing cells in different growth phases, including Lag phase (20% growth area occupied, 1 : 20 split), Log phase (75% growth area occupied, 1 : 10 split) and plateau phase (>95% growth area occupied, 1 : 2 split; Supplementary Information, Figure 1). From those T75 flasks, equal numbers of cells were transferred to 6well plates to perform the drug treatment. First, drug treatment after homogeneous subculture was performed by seeding 250’000 cells exclusively from the Log phase into six wells of a 6-well plate (Sarstedt GmbH, Nümbrecht, Germany). Cells were counted with a MOXI Z Mini automated cell counter (Orflo Technologies, Ketchum, ID, USA). Additionally, the treatment after heterogeneous subculture was performed by seeding 250’000 cells from the Lag phase, from the Log phase and from the plateau phase each into two wells of a 6-well plate (Figure 1). Each well contained 250’000 cells in 2 mL complete medium. After seeding, SW480 cancer cells were left to adhere for approximately 21 h. Then, medium was exchanged with solvent control medium or medium containing arsenic trioxide (ATO; Sigma–Aldrich, USA) and incubated for 24 h. ATO stock solutions were prepared according to supplier instructions (pH 7.4). The final ATO concentration (5 μM) was used as previously reported by us.^31^ Cell growth^39^ and motility^40^ was continuously monitored during adhesion and treatments using a Cellwatcher M and the software PHIOme (Version 2022, PHIO scientific GmbH, Munich, Germany).

#### Sample work-up

All steps for the sample collection process were performed on ice. The supernatant of each well (2 mL) was transferred into a 15 mL Falcon® tube, centrifuged (1233 g, 5 min, 4 °C) to remove cellular debris and transferred again into a 15 mL Falcon® tube containing ice-cold absolute ethanol (8 mL) and 5 μL of an internal eicosanoid standard mixture (12S-hydroxyeicosatetraenoic acid (HETE)-d8, 15SHETE-d8, 20-HETE-d6, 5-oxo-eicosatetraenoic acid (ETE)-d7, prostaglandin E2 (PGE2)-d4 and 11,12-dihydroxy-5Z,8Z,14Z-eicosatrienoic acid (DiHETrE)-d11 (Cayman Chemical, Tallinn, Estonia)). The concentrations of the internal standards can be found in Supplementary Information, Table 1. Afterwards, the tube was closed, inverted once and sealed with parafilm. The samples were stored overnight at –20°C and then processed according to the oxylipin and fatty acid analysis protocol. The adherent SW480 cells were washed twice with cold Tris-buffered saline (TBS; 25 mM Tris·HCl, 150 mM NaCl, pH 7.4) and then, the wash buffer was completely removed. A volume of 80 μL of 4% [wt/vol] sodium deoxycholate lysis buffer (SDC; 0.4 g SDC, 500 μL Tris·HCl (pH 8.8) in 9.5 mL H_2_ O) was added to the well. The cells were scraped off and the whole cell lysate (WCL) was transferred to an Eppendorf® tube. This step was repeated with 50 μL SDC and the remaining WCL was also added to the corresponding tube, which was subsequently heated to 95 °C for 5 min. These samples were stored at –20 °C until (phospho-)proteomic processing.

### Oxylipin and fatty acid analysis

#### Sample Preparation

The analysis of oxylipins and fatty acids was performed as previously described.^4^ The eicosanoid samples in Flacon® tubes were centrifuged (30 min, 4536 g, 4 °C) and the clear supernatants were transferred into new 15 mL Falcon® tubes. EtOH was evaporated *via* vacuum centrifugation at 37 °C until the original sample volume (2 mL) was restored. Samples were loaded onto preconditioned StrataX solid-phase extraction (SPE) columns (30 mg mL^−1^, Phenomenex, Torrance, CA, USA) using Pasteur pipettes. SPE columns were washed with icecold H_2_ O (5 mL, MS grade, VWR International, Vienna, Austria) and elution of analytes was carried out with ice-cold MeOH (500 μL, MS grade, VWR International, Vienna, Austria) containing 2% formic acid (FA; ≥99%. VWR International, Vienna, Austria). The eluted samples were dried using a gentle nitrogen stream at room temperature. The dried samples were reconstituted in 150 μL reconstitution buffer (H_2_O : ACN : MeOH [vol% 65 : 31.5 : 3.5] + 0.2% FA). Then, the samples were directly transferred into an autosampler held at 4 °C and subsequently analyzed *via* LC-MS/MS.

#### LC–MS/MS Analysis

For the LC-MS analyses, separation of analytes was achieved using a Vanquish^™^ UHPLC system (Thermo Fisher Scientific^™^, Vienna, Austria) equipped with a reversed-phase Kinetex^®^ C18 column (2.6 μm XB-C18, 100 Å, LC column 150 × 2.1 mm, Phenomenex, Torrance, CA, USA). Flow rate was set to 200 μL·min^−1^, the injection volume was 20 μL and all samples were measured in technical duplicates. The LC column oven was set to 40 °C and the autosampler was set to 4 °C. A gradient flow profile was applied (mobile phase A: H_2_O + 0.2% FA, mobile phase B: ACN : MeOH [vol% 90 : 10] + 0.2% FA) starting at 35% B and increasing to 90% B (1–10 min). After further increasing to 99% B within 0.5 min and keeping it for 5 min, solvent B was decreased to 35% B within 0.5 min and held for 4 min to equilibrate the column, resulting in a total run time of 20 min. The Vanquish^™^ UHPLC system was hyphenated to a high-resolution quadrupole orbitrap mass spectrometer (Thermo Fisher Scientific^™^ QExactive^™^ HF hybrid quadrupole orbitrap mass spectrometer), which was equipped with a HESI source operating in negative ionization mode. Spray voltage was set to 3.5 kV, capillary temperature was set to 253 °C, sheath gas and auxiliary gas were set to 46 and 10 arbitrary units, respectively. Data were recorded in the scan range of *m/z* 250–700 on the MS1 level with a resolution of 60’000 (*m/z* 200). A Top2 method with a resolution of 15’000 (*m/z* 200) was selected using HCD fragmentation with a normalized collision energy of 24 for data-dependent acquisition. Additionally, an inclusion list was generated with 33 precursor masses, which are specific for oxylipins and their precursor fatty acids (Supplementary Information, Table 2).

#### LC–MS/MS Data Processing

For the data analysis, raw files generated by the QExactive^™^ HF hybrid quadrupole orbitrap high-resolution mass spectrometer were checked manually using the Xcalibur^™^ Qual Browser software (version 4.1.31.9, Thermo Fisher Scientific^™^, Bremen, Germany) by comparing reference spectra from the LIPIDMAPS depository library from July 2020.^75^ In general, identification of analytes was performed using exact mass, retention time and MS/MS fragmentation pattern. For relative quantification, the TraceFinder software (version 4.1, Thermo Fisher Scientific^™^, Bremen, Germany) was applied, allowing a mass deviation of 5 ppm. The resulting data containing peak areas of each analyte were loaded into the R software package environment (version 4.2.0)^76^ and log_2_-transformed. For normalization, the log_2_transformed mean peak area of the internal standards was subtracted from the log_2_-transformed analyte areas to correct for variances arising from sample extraction and LC-MS/MS analysis. Then, each log_2_ -transformed area was increased by adding (x+20) to obtain a similar value distribution compared to label-free quantification in proteomics. Missing values were imputed using the minProb function of the imputeLCMD package (version 2.1).^77^ Volcano plots were produced using the ggplot2 package (version 3.4.0).^78^ In an unpaired T-test, p-values <0.05 were considered to indicate statistical significance and the BenjaminiHochberg procedure was applied as multiple testing correction to all p-values.

### Proteomics and Phosphoproteomics

#### Sample Preparation

Proteomic and phosphoproteomic samples were prepared using a modified version of a previously described protocol^79^ and employing an adapted version of the EasyPhos platform.^80^ WCLs were thawed and lysed using the S220 Focused-ultrasonicator (Covaris, LLC., Woburn, MA, USA). Protein concentrations were determined *via* bicinchoninic acid assay (BCA)-assay. Between 130–220 μg of protein was reduced and alkylated with tris(2-carboxyethyl)phosphine (TCEP) and 2-chloroacetamide (2CAM) for 5 min at 45 °C, followed by 18 h digestion with Trypsin/Lys-C (1:100 enzyme-to-substrate ratio) at 37 °C. An aliquot of 20 μg peptide was taken from each sample for global proteome analysis and dried in a vacuum concentrator. Then, the samples were reconstituted in styrenedivinylbenzene-reverse phase sulfonate (SDB-RPS) loading buffer (99% iPrOH, 1% TFA) and desalted *via* SDB-RPS StageTips. Desalted global proteome samples were reconstituted in 5 μL formic acid (30%) containing synthetic standard peptides at 10 fmol and diluted with 40 μL loading solvent (98% H_2_O, 2% ACN, 0.05% TFA). For phosphopeptide enrichment, the digested samples were mixed with enrichment buffer (52% H_2_O, 48% TFA, 8 mM KH_2_PO_4_) and incubated with TiO_2_Titansphere beads (GL Sciences, Japan), followed by sample clean-up *via* C_8_StageTips. Phosphopeptides were eluted and subsequently dried in a vacuum concentrator. Phosphopeptides were reconstituted in 15 μL MS loading buffer (97.7% H_2_O, 2% ACN, 0.3% TFA).

#### LC-MS/MS analyses

LC-MS/MS analyses were performed employing a timsTOF Pro mass spectrometer (Bruker Daltonics, Bremen, Germany) hyphenated with a Dionex UltiMate^™^3000 RSLCnano system (Thermo Scientific, Bremen, Germany). Samples were analyzed in data-dependent acquisition mode by label free quantification (LFQ) shotgun proteomics similarly to a recently published method.^79^ The injection volume was 10 μL and 2 μL for analyzing phosphoproteomes and global proteomes, respectively. Samples were loaded on an Acclaim^™^ PepMap^™^C18 HPLC pre-column (2 cm × 100 μm, 100 Å, Thermo Fisher Scientific^™^, Vienna, Austria) at a flow rate of 10 μL min^− 1^ MS loading buffer. After trapping, peptides were eluted at a flow rate of 300 nL min^− 1^ and separated on an Aurora series CSI UHPLC emitter column (25 cm × 75 μm, 1.6 μm C18, Ionopticks, Fitzroy, Australia) applying a gradient of 8–40% mobile phase B (79.9% ACN, 20% H _2_ O, 0.1% FA) in mobile phase A (99.9% H _2_ O, 0.1% FA) over 85 min.

#### Data analysis

Protein identification was performed *via* MaxQuant^81^ (version 1.6.17.0) employing the Andromeda search engine against the UniProt Database^82^ (version 11/2021, 20’375 entries). Search parameters were set as previously described.^79^ A mass tolerance of 20 ppm for MS spectra and 40 ppm for MS/MS spectra, a PSM–, protein– and site-false discovery rate (FDR) of 0.01 and a maximum of two missed cleavages per peptide were allowed. Matchbetween-runs was enabled with a matching time window of 0.7 min and an alignment time window of 20 min. Oxidation of methionine, N-terminal protein acetylation and phosphorylation of serine, threonine and tyrosine were set as variable modifications. Carbamidomethylation of cysteine was set as fixed modification. Global proteome data analysis was performed *via* Perseus (version 1.6.14.0). Proteins with at least 60% quantification rate in at least one group were considered for analysis. A two-sided Student’s T-test with S0 = 0.1 and a FDR cut-off of 0.05 was applied for identifying multiple testing-corrected significantly regulated proteins. Phosphoproteomic data processing and analysis was performed according to a modified version of the PhosR protocol^83^ employing the R-packages PhosR^47^ (version 1.4.0), limma^69^ (version 3.50.3) and dplyr^84^ (version 1.0.10). Highest intensity values of phosphosites identified in multiple phosphopeptide isoforms were used for analysis. Reverse matches and potential contaminants were filtered. Only class I phosphosites with localization probability ≥0.75 with at least 60% quantification rate in at least one group were considered. Tail-based imputation of missing values was performed and data was centered across their median. Differentially phosphorylated sites between conditions were determined, phosphosites with an adjusted pvalue ≤0.05 were considered as significantly regulated. For kinase-substrate mapping, an analysis of variance (ANOVA) test was performed and phosphosites with an adjusted p-value <0.05 and with difference >0 were submitted to the kinaseSubstrateScore function of PhosR with default parameter settings (numMotif = 5, numSub = 1).

#### Data visualization

If not otherwise specified, data visualization was performed in RStudio^76^ (version 2021.09.0) employing the packages ggplot2^78^ (version 3.3.6), ggpubr^85^ (version 0.4.0), ggrepel (version 0.9.1)^86^ and VennDiagram (version 1.7.3).^87^

#### CausalPath analysis

CausalPath^7^ was utilized to analyze causal networks integrating global proteomics and phosphoproteomics using pre-processed LFQ intensity values and standard parameter settings: value transformation = significant-change-of-mean and FDR threshold = 0.1 for protein and phosphoprotein. The resulting networks were visualized in Newt.^88^

## Statistical analysis

Normalized oxylipin and fatty acid data was statistically analyzed using an unpaired T-test (p-values <0.05) with limma. P-values were multiple-testing corrected by the Benjamini-Hochberg procedure.

Proteomics data was analyzed in Perseus (Version 1.6.14.0) using a two-sided T-test with S0 = 0.1 and a permutation-based FDR cut-off of 0.05 was applied for identifying multiple-testing corrected significantly regulated proteins.

Differentially phosphorylated sites between conditions were determined. Phosphosites with an adjusted p-value ≤0.05 were considered as significantly regulated. For kinase-substrate mapping, an analysis of variance (ANOVA) test was performed and phosphosites with an adjusted p-value <0.05 and with difference >0 were submitted to the kinaseSubstrateScore function of PhosR with default parameter settings (numMotif = 5, numSub = 1).

## Data availability

All (phospho-)proteomics data was submitted to the ProteomeXchange Consortium (http://proteomecen-tral.proteomexchange.org) and is available in the PRIDE partner repository^89^ with the dataset identifier PXD039768.

## Acknowledgements

The authors are grateful to the Core Facility of Mass Spectrometry at the Faculty of Chemistry and the Joint Metabolome Facility, members of the Vienna Life-Science Instruments (VLSI). Figures were created and assembled using BioRender.com.

## Author contributions

LS performed cell culture, sample workup and Cellwatcher acquisition. PB performed proteomics and phosphoproteomics. GH performed oxylipin and fatty acid analysis. AB, GH and PB acquired and processed data. VP, LG and TM processed data. PB assembled figures. CG, SMM and PP planned and supervised the study. PB, GH, LS, VP, CG and SMM wrote the initial draft of the manuscript, which was edited and approved by all authors.

## ADDITIONAL INFORMATION

Supporting Information to this article is available. This includes a supplementary information file containing figures and tables (doc). Additionally, a high-resolution file of Figure 06 (tif) and two supplementary tables including lists of significant regulations (exe) can be accessed.

