## Supplementary Information for "Multi-Omics Data of Perturbation Studies are Determined by Memory Effects from Subculture"

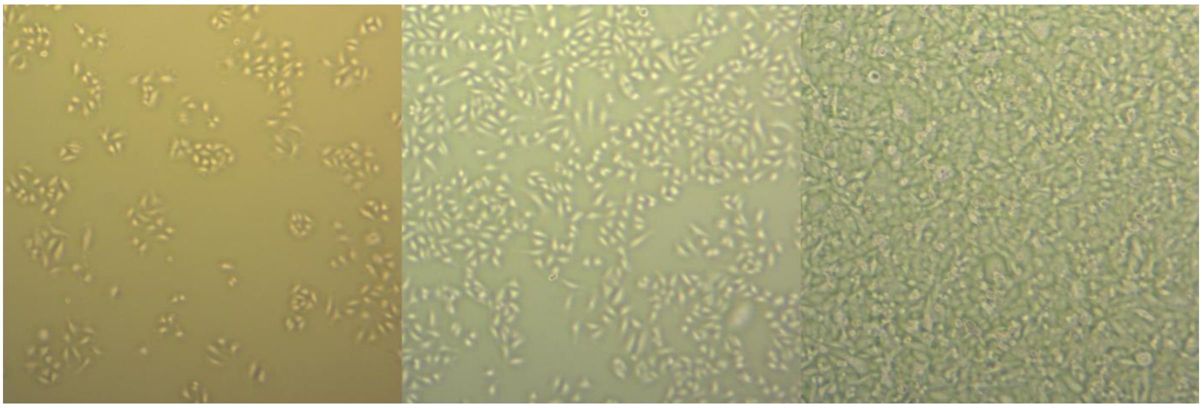

**Supplementary Information, Figure 1:** SW480 cancer cells in different growth phases used for subculture: Images of cells 48 h after seeding in the Lag phase (*left*, 20% growth area occupied, 1:20 split), in the Log phase (*middle*, 75% growth area occupied, 1:10 split) and in the plateau phase (*right*, >95% growth area occupied – confluence, 1:2 split). Images were obtained at 4× magnification using a light microscope (Zeiss).

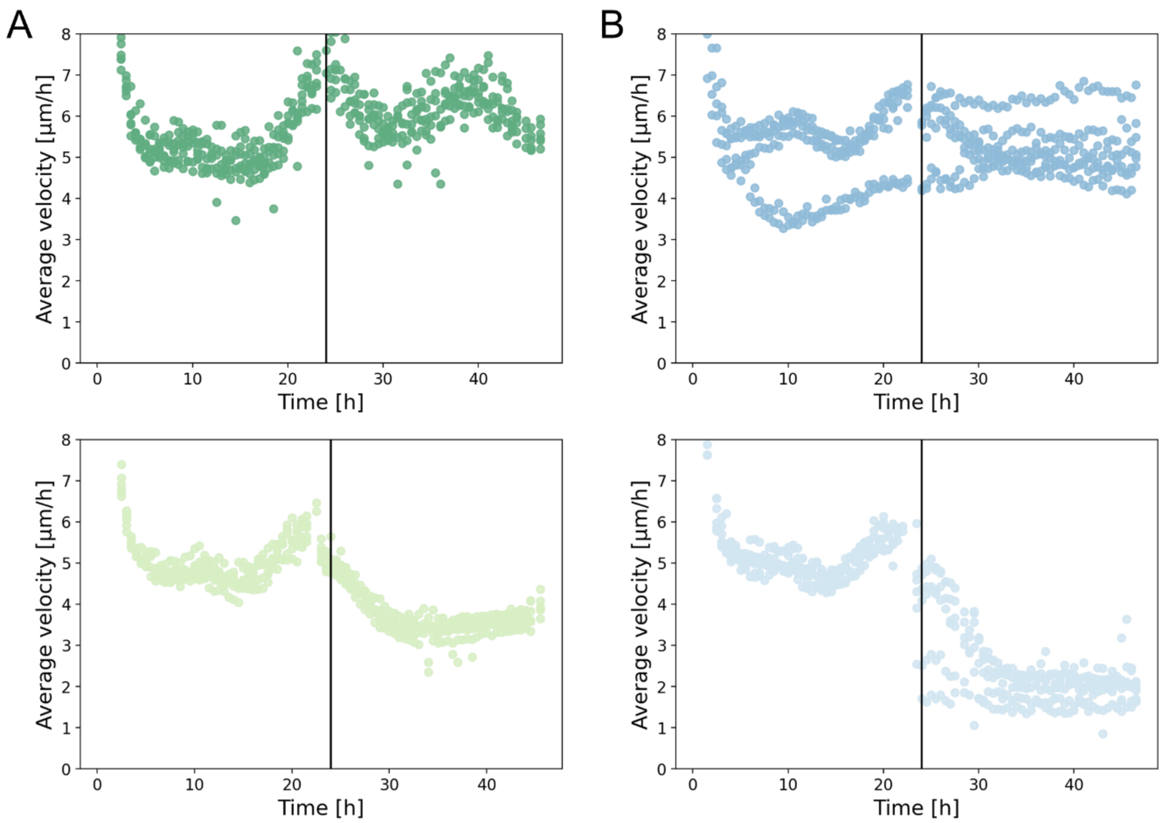


**Supplementary Information, Figure 2:** Cell motility of samples in individual wells was followed throughout the adhesion and treatment phases after subculture. Medium was replaced after one day either with vehicle control or arsenic trioxide (5 µM) and incubated for further 24 h. Treatment with arsenic trioxide similarly inhibited cell motility after homogeneous (**A**, green) and heterogeneous (**B**, blue) subcultures.

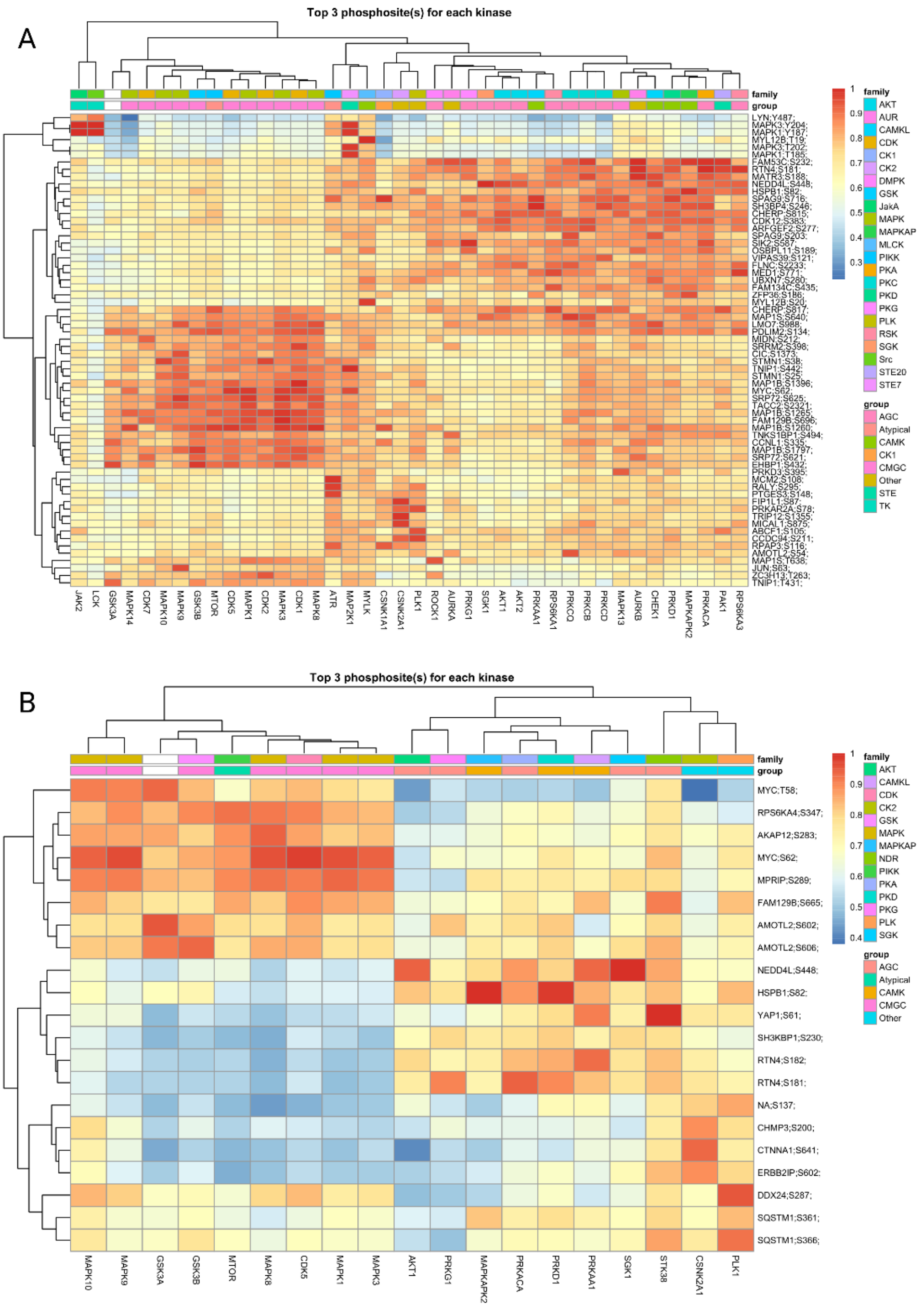


**Supplementary Information, Figure 3:** Heatmap of the combined scores retrieved for the top 3 phosphosites (highest combined scores) of all evaluated kinases. (**A**) Significantly upregulated phosphosites after homogeneous subculture resulted in the annotation of 40 kinases to their corresponding top 3 scoring phosphosites. (**B**) Significantly upregulated phosphosites after heterogenous subculture resulted in the annotation of 19 kinases to their corresponding top 3 scoring phosphosites.

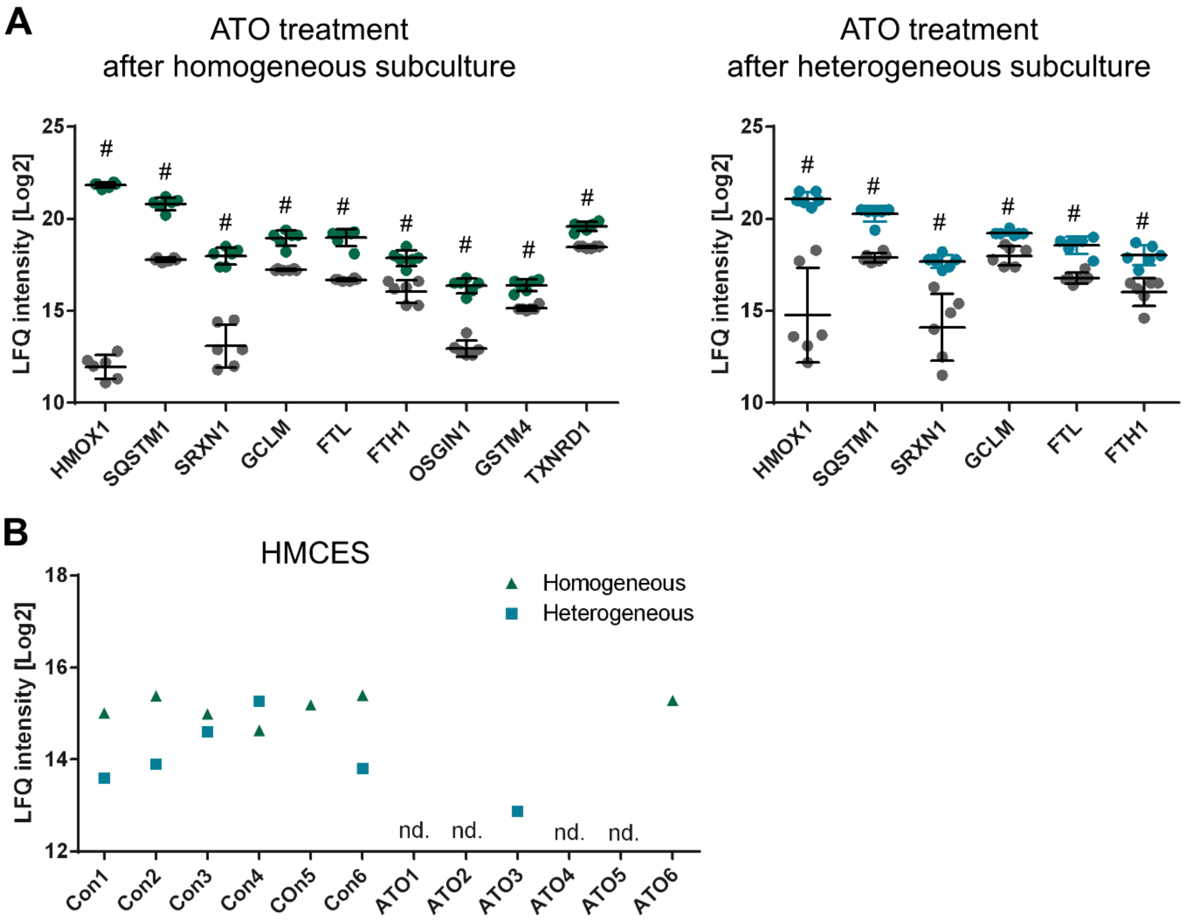


**Supplementary Information, Figure 4:** Additional perturbations that were not detected in the perturbation network, and only after manual data interpretation. (**A**) Induction of Nrf2 target genes in SW480 cancer cells after treatment with arsenic trioxide (ATO) after homogeneous (*left*) and heterogeneous subculture (*right*). Controls are shown in grey and ATO-treated cells are shown in green or blue. (**B**) DNA damage may be reduced by the suicide enzyme HMCES, which was stably expressed in controls, but largely consumed after ATO-treatment. nd. = not detected.

**Supplementary Information, Table 1:** List of internal eicosanoid standards and their respective concentration in each sample.

| Internal standard | Abbreviation | c [pg/µL] |
|---|---|---|
| 12S-hydroxyeicosatetraenoic acid-d8 | 12S-HETE-d8 | 6.67 |
| 15S-hydroxyeicosatetraenoic acid-d8 | 15S-HETE-d8 | 6.67 |
| 5-oxo-eicosatetraenoic acid-d7 | 5-OxoETE-d7 | 20.00 |
| 11,12-dihydroxy-5Z,8Z,14Z-eicosatrienoic acid-d11 | 11,12-DiHETrE-d11 | 6.67 |
| prostaglandin E2-d4 | PGE2-d4 | 13.33 |
| 20-hydroxyeicosatetraenoic acid-d6 | 20-HETE-d6 | 6.67 |

**Supplementary Information, Table 2:** Inclusion list of 33 oxylipins and their precursors used for mass spectrometric analysis.

| Mass [m/z] | Mass [m/z] | Mass [m/z] | Mass [m/z] |
|---|---|---|---|
| 254.2245 | 311.2228 | 333.2071 | 359.2222 |
| 275.2011 | 313.2384 | 335.2222 | 367.3576 |
| 277.2167 | 315.1966 | 337.2384 | 375.2171 |
| 279.2324 | 317.2122 | 343.2279 | |
| 281.2480 | 319.2279 | 348.3069 | |
| 283.2637 | 321.2435 | 349.2020 | |
| 293.2122 | 325.2382 | 351.2177 | |
| 295.2279 | 327.2324 | 353.2328 | |
| 301.2168 | 327.2781 | 355.2428 | |
| 303.2324 | 329.2480 | 357.2585 | |
